# LGFN: A Dynamic Gating Framework for Lyrics-Audio Alignment in Music Emotion Recognition

**DOI:** 10.1101/2025.08.11.669801

**Authors:** Yinghui Wang, Mengqi Zang, Huasen Zhang, Xin Shan, Xinyue Wang

**Affiliations:** The Graduate School Arts and Culture, Sangmyung University, Seoul, South Korea

## Abstract

Music emotion recognition (MER) is challenging due to the ambiguous nature of audio features. For example, a fast tempo might indicate excitement or anger. To address this issue, we propose the Lyrics-Aware Gate Fusion Network LGFN, a novel cross-modal dynamic fusion approach. This network extracts audio features using ResNet for Mel spectrograms and 1D CNN for waveforms, while leveraging BERT for lyric embeddings. The key innovation of LGFN lies in its attentional alignment layer AAL, which effectively bridges the temporal misalignment between lyrics and audio. Additionally, the dynamic gate fusion module DGFM adaptively adjusts the weight of audio and text features based on their reliability. This allows the model to automatically determine the contribution of each modality, enhancing the accuracy of emotion recognition. Ablation studies on the music emotion recognition datasets MER1101 and DEAM2015 demonstrate the effectiveness of our proposed AAL and DGFM, while visualizing the gate weights offers insights into the model’s decision-making process. We also conduct comprehensive experiments on different human multimodal emotion recognition benchmarks, which show the superiority of our approach over the state-of-the-arts.

## Introduction

Music emotion recognition (MER) is an important area of research that enables computers to accurately comprehend human emotions and provide intelligent responses to meet human requirements. With the rising demand for music consumption and the explosive growth of music content, MER demonstrates its critical position in music understanding and applications. It has been widely used in personalized music recommendation [1], music therapy [2], music education [3], music generation [4], etc. In educational settings, it can monitor students’ emotional states, helping educators understand their academic progress and overall condition. In healthcare, it assists doctors in grasping patients’ emotional states for personalized medical services. It also enhances intelligent customer service systems by detecting users’ emotional needs and offering more personalized services. However, the complexity of music emotions makes accurate recognition challenging.

Traditional single-modal approaches in music emotion recognition are limited as they rely solely on audio features or lyrics, failing to capture the full spectrum of emotional cues [5]. Audio features alone cannot convey the semantic depth of a song, while lyrics lack crucial acoustic context. To address this, researchers have explored two main models for emotion prediction: the discrete emotion model [6, 7], which uses categorical adjectives like happiness and anger, and the dimensional emotion model [8–11], which maps emotions onto a continuous scale. Russell’s two-dimensional valence-arousal (V-A) model [12], in particular, transforms emotion prediction into a two-dimensional regression problem. This paper focuses on Dynamic Music Emotion Recognition (DMER), predicting music emotion using continuous V-A values at short intervals. Meanwhile, multi-modal fusion methods offer a more comprehensive solution by integrating audio and lyrical data [13, 14]. However, practical challenges arise from the asynchrony of multimodal streams due to varying sampling rates. Previous works [15–18] have used predefined word-level alignment to address this, but the process is labor-intensive and requires domain knowledge.

Recent advances in deep learning have led to the development of more sophisticated multimodal fusion techniques. For instance, Tsai et al. proposed the Multimodal Transformer (MulT) approach [19], which uses modality reinforcement units to fuse crossmodal information from unaligned data sequences. By learning directional pairwise attention between elements across modalities, MulT can implement multimodal fusion from asynchronous sequences without explicit data alignment [20]. Despite its advantages, the MulT approach reinforces each modality pair independently, without exchanging information between them. This limitation means that multimodal fusion only occurs between each directional modality pair, rather than across all involved modalities. Furthermore, the independent pairwise fusion approach fails to fully exploit the high-level features of the source modality. Each directional modality pair reinforces the target modality by attending to the low-level features of the source modality. This semi-shallow structure limits the exploration of deep interactions across modalities.

To address these limitations, researchers have explored various fusion strategies, including early fusion, late fusion, and hybrid fusion methods. Early fusion involves combining data from different modalities at the input stage [21–24], while late fusion combines high-level features extracted from each modality [24–26]. Hybrid fusion combines both early and late fusion approaches to leverage their respective advantages [27, 28]. Additionally, intermediate layer fusion methods perform fusion within the deep network’s layers, allowing for more flexible and effective integration of multimodal information. Among the existing studies, Long Short-Term Memory (LSTM) has received extensive attention in DMER due to its superiority in sequence modeling [8, 29–31]. Convolutional Neural Network (CNN) is used to extract features in many fields. Researchers have recently focused on improving emotion recognition accuracy using a combination of CNN and Recurrent Neural Network (RNN) [9, 32–34]. However, LSTM-based models still use handcrafted features as input, and some widely used handcrafted feature operations will lose high-level features. The CNN-RNN based model mainly uses a fixed-scale CNN. Due to its fixed receptive field, the learned CNN features are limited, and the emotional crucial features of different fields of view are not extracted. Moreover, various problems exist in existing music emotion datasets, which also hinder the progress of DMER.

In this paper, we propose the Lyrics-Aware Gate Fusion Network LGFN, a novel cross-modal dynamic fusion approach for music emotion recognition. LGFN combines ResNet and 1D CNN for audio feature extraction from Mel spectrograms and waveforms, respectively, while utilizing BERT [35] for deep lyrical embeddings. The network’s architecture is distinguished by two core components: an attentional alignment layer and a dynamic gate fusion module. The attentional alignment layer corrects temporal misalignment by synchronizing lyrical and audio features, ensuring accurate correlation of emotional cues. The dynamic gate fusion module adaptively weights the contributions of audio and lyrical features based on their reliability, allowing the model to focus on the most informative modality for each prediction.

To sum up, the main contributions of this work can be summarized as follows:

- We propose a cross-modal attentional alignment layer that resolves temporal misalignment between lyrics and audio streams through positional encoding, effectively bridging asynchronous multimodal sequences.
- We introduce a self-adaptive gating mechanism that dynamically modulates audio-text feature contributions based on semantic reliability, enhancing emotion discernment while providing model interpretability via visualized gate weights.
- Our approach demonstrates superior performance compared to existing state-of-the-art methods across various music emotion recognition benchmarks.

## Related Work

This section briefly reviews related works in two categories: uni-modal emotion recognition and multi-modal emotion recognition.

### uni-modal emotion recognition

Emotion recognition has been extensively studied using single modalities, including text, audio, and visual information. In text-based emotion recognition, traditional methods often rely on the bag-of-words model [36, 37], which, despite its simplicity, fails to capture word order and contextual information. To overcome these limitations, word embedding techniques such as word2vec [38] and GloVe [39] have been introduced. These models learn semantic representations of words using neural networks but still lack the ability to capture context-specific meanings. More recent approaches generate context-aware embeddings by considering both left and right contexts within a sentence. Additionally, sentiment-specific embeddings have been developed to better capture emotional semantics at the word level.

For facial emotion recognition, traditional methods relied on handcrafted features like Local Binary Pattern (LBP) [40] and Histograms of Oriented Gradient (HOG) [41]. These methods require manual feature extraction and often overlook significant semantic information. With the rise of deep learning, CNNs have become the dominant approach. They automatically learn optimal features from facial images and have been applied in various tasks such as action recognition and dynamic scene recognition.

### multi-modal emotion recognition

Multimodal emotion recognition integrates information from various modalities to enhance the detection of emotional states [42, 43]. Early approaches, such as early and late fusion strategies combined with probabilistic graphical models, have been traditionally employed. However, these methods often neglected the inherent dependencies between elements of sequences from different modalities, which are essential for effective multimodal fusion. To address this, recent works have introduced sophisticated methods like manual alignment of visual and acoustic sequences with textual words before training, and the use of hierarchical attention mechanisms and cyclic translation for multimodal fusion.

To tackle the challenges of unaligned multimodal sequences, crossmodal attention mechanisms have been developed. These mechanisms learn inherent correlations across modalities by reinforcing one modality with information from others through directional pairwise attention [19, 44]. Various deep learning-based fusion methods have been categorized into model-agnostic and intermediate layer fusion. Model-agnostic methods perform fusion outside of a specific deep learning model, while intermediate layer fusion integrates the fusion process within the deep network’s layers. These approaches aim to leverage the complementary strengths of different modalities to achieve more accurate emotion recognition than single-modality approaches.

### Methodology

#### Task Definition

In this work, corresponding to the lyrics and audio of music, the music emotion recognition task in volves two major modalities, i.e., Text (T) and audio (A). Denote the input sequences from the corresponding modalities by 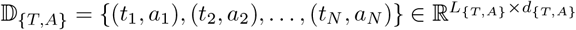, where (*t*_*i*_, *a*_*i*_) represents the *i*^*th*^ music clip pair in the music, *N* is the number of music clip in the dialogue. *L*_(·)_ and *d*_(·)_ represent the sequence length and feature dimension, respectively. Musci emotion recognition task requires identifying the emotion category labels *E* = {*e*_1_, *e*_2_, …, *e*_*N*_} of the entire dialogue (*t*_1_, *a*_1_), (*t*_2_, *a*_2_), …, (*t*_*N*_, *a*_*N*_). The objective of this study is to achieve efficient cross-modal dynamic fusion from multimodal data sequences, thereby enhancing the accuracy and robustness of music emotion recognition.

#### Model overview

The architecture of our proposed model LGFN for Music Emotion Recognition (MER) is depicted in Fig 1 and primarily consists of three main components: the audio-lyric feature extraction part, the cross-modal fusion part, and the emotion recognition part. Specifically, Audio features are extracted using ResNet for Mel spectrograms and 1D CNN for waveforms, while lyric features are generated using BERT embeddings. In addition, cross-modal fusion is achieved via an Attentional Alignment Layer (AAL) to correct temporal misalignment and a Dynamic Gate Fusion Module (DGFM) to weigh modality contributions adaptively. Finally, we obtain the recognition results by designing classifiers for the MER tasks in the emotion recognition part.

**Fig 1.**
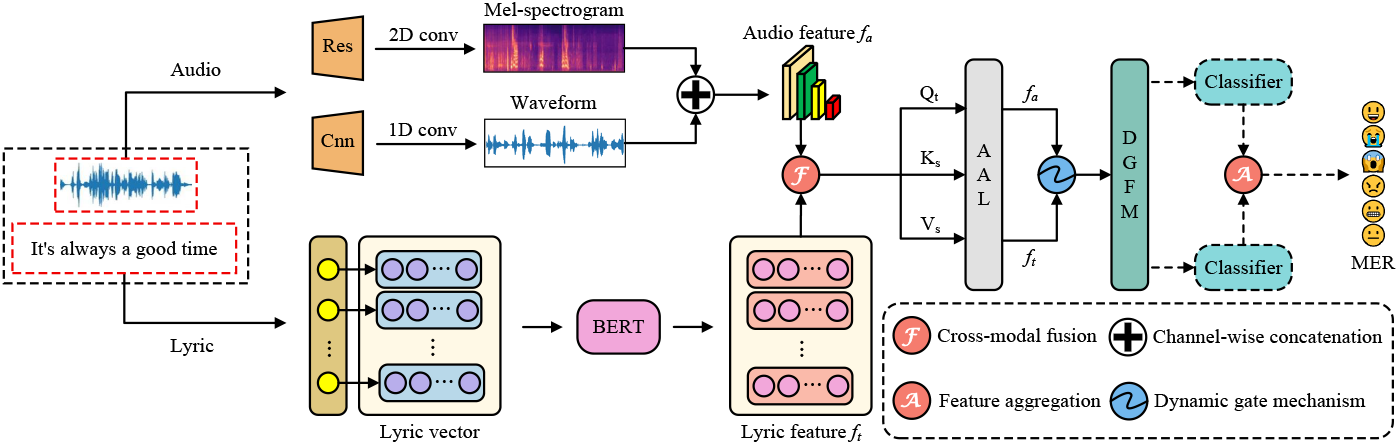
The general overview of the proposed LGFN architecture. The input music is subjected to feature extraction by both the audio branch and the lyrics branch for music emotion recognition. The AAL denote Attention alignment layer and the DGFM denote Dynamic gate fusion module.

The model, trained in an end-to-end manner, processes input sequences from different modalities using respective feature extraction techniques. Fig 2 displays the information flow across the attention alignment layers. Denote the processed sequences from the audio and lyric modalities as 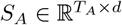 and 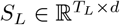, respectively. The common message *S*_*C*_ is initialized by concatenating the low-level sequences: *S*_*C*_ = [*S*_*A*_, *S*_*L*_], where 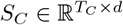 and *T*_*C*_ = *T*_*A*_ + *T*_*L*_. The AAL corrects temporal misalignment, and the DGFM adjusts modality weights. The reinforced features 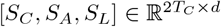 are passed through a transformer layer, followed by fully connected layers to classify emotion labels.

**Fig 2.**
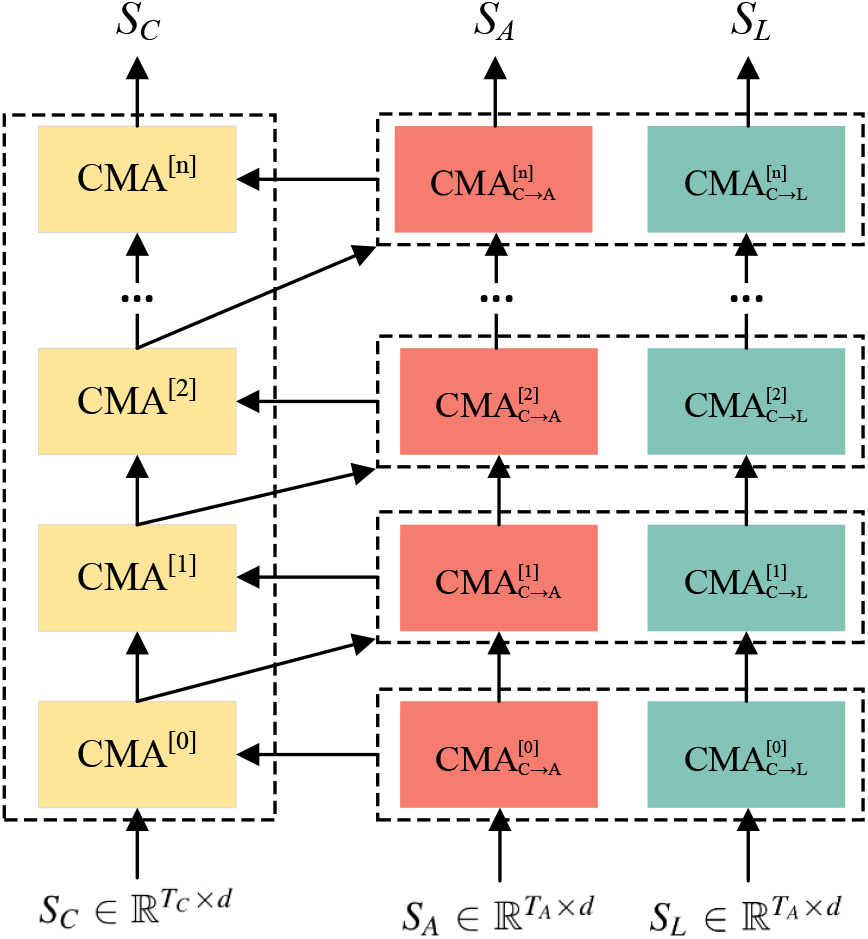
The information flow across the attention alignment layer of the proposed model. *CMA*^[*i*]^ represents the common feature which is operated by the cross-modal attention from the next layer. 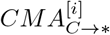 denotes the modality alignment unit in which the corresponding target modality is reinforced by the common message

### Feature Extraction

#### Attentional Alignment Layer

To address the temporal misalignment between lyrics and audio features, we design the Attentional Alignment Layer AAL. This layer employs crossmodal attention mechanisms to synchronize the features extracted from both modalities, ensuring that the emotional cues are accurately correlated.

In the AAL, we utilize a crossmodal attention operation that reinforces the target modality with information from a source modality by learning the directional pairwise attention between them. Let 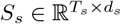 denote the data sequence from the source modality and 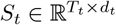 denote the data sequence from the target modality, where *s, t* ∈ {*T, A*}. The crossmodal attention unit involves Queries, Keys, and Values, defined as follows:

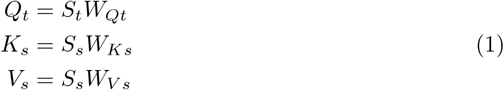

Where 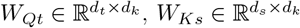, and 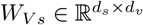 are learnable weight matrices. One individual head of crossmodal attention is defined as:

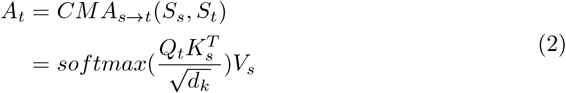

Where 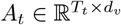. The full crossmodal attention operation with *h* heads is represented as 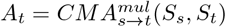, where 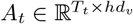.

The target modality is reinforced by encouraging the model to attend to crossmodal interaction between elements, allowing the network to focus on the most relevant features for emotion recognition. This crossmodal attention mechanism effectively bridges the temporal gap between lyrics and audio, enhancing the model’s ability to capture synchronized emotional cues from both modalities.

#### Dynamic Gate Fusion Module

The Dynamic Gate Fusion Module (DGFM) is designed to resolves modality reliability fluctuations in cross-modal music emotion recognition by adaptively weighting contributions of aligned audio and lyric features. This self-gating mechanism computes a dynamic fusion weight *g* ∈ [0, 1] through learnable projections of the input features:

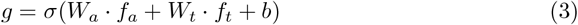

where *f*_*a*_ and *f*_*t*_ represent the aligned audio and lyric features, *W*_*a*_ and *W*_*t*_ are learnable weight matrices, *b* is a bias term, and *σ* is the sigmoid activation function. The fused representation *f* ∈ ℝ is then derived as:

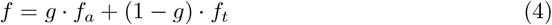

where the fusion weight *g* prioritizes audio during acoustically salient segments, while 1 − *g* amplifies lyrics in semantically rich contexts. The final fused feature *f* is a weighted sum of the audio and lyric features, capturing the most informative cues for emotion recognition.

### Experiments

#### Datasets

We conduct our experiments on the IEMOCAP dataset [45], DEAM2015 dataset [46] and the dataset MER1101 [47]. The details of each dataset are given below.

#### IEMOCAP

This dataset [45] comprises 4,453 video clips with a predefined partition: 2,717 training samples, 798 validation samples, and 938 testing samples. Acoustic and visual features were extracted at sampling rates of 12.5 Hz and 15 Hz, respectively.

Following prior methodology [48], we focus on recognizing four emotion categories—happy, sad, angry, and neutral—in each clip. The task is established as multi-label classification since sad and angry emotions may coexist within a single clip. **DEAM** Developed for the MediaEval Emotion in Music task, the DEAM2015 benchmark [46] contains 431 thirty-second training samples and 58 full-length evaluation songs. As the predominant resource for dynamic music emotion recognition, its evaluation set demonstrates notably low valence reliability with Cronbach’s *α* = 0.29 ± 0.94. Significant performance discrepancies emerge between training and evaluation sets, particularly in valence dimensions, attributable to spatiotemporal variations and annotator inconsistencies during emotion labeling.

#### MER1101

Similarly based on Russell’s valence-arousal model, MER1101 [47] comprises 1,101 web-sourced music clips spanning 16.5 to 125.5 seconds, featuring both discrete and dimensional emotion labels. Each track received annotations from three experts and ten students through a rigorous protocol: initial emotion adjective labeling followed by valence-arousal scoring after song familiarization. Student annotations achieved Cronbach’s *α* values of 0.6295 ± 0.3574 for arousal and 0.5624 ± 0.3766 for valence, while expert annotations yielded 0.3556 ± 0.3442 for arousal and 0.2420 ± 0.3148 for valence.

### Evaluation Metrics

We use the Concordance Correlation Coefficient (CCC), Pearson Correlation Coefficient (PCC), and Root Mean Square Error (RMSE) as evaluation metrics. Each metric is calculated using the ground-truth and predicted V-A values for each song, with the results averaged across all songs. The CCC combines the characteristics of PCC and RMSE, making it suitable for assessing both the trend of emotional changes and the disparity between predictions and ground-truth. Therefore, we consider CCC to be the primary evaluation metric.

For the IEMOCAP dataset, we use accuracy and F1-score to evaluate the models. Additionally, in line with [48], we employ standard binary F1 rather than the weighted version.

### Implementation details

For DEAM2015, we strictly followed the predefined training/evaluation set configuration. Regarding MER1101, 925 tracks exceeding 30-second duration were selected and randomly divided into training and evaluation sets. Training tracks were segmented into 30-second clips yielding 1,526 samples, while the evaluation set retained 185 full-length songs. To ensure statistical reliability, 5-fold cross-validation was implemented on both datasets. Log Mel-spectrograms were extracted using the librosa toolbox with 128 Mel bands at 44.1 kHz sampling rate, employing 60 ms windows and 10 ms hop sizes. Model training utilized the Adam optimizer over 100 epochs with batch size 32, incorporating early stopping to prevent overfitting. Arousal prediction employed C-CC loss, while valence used RMSE loss.

The hyperparameters for each benchmark are shown in Table 1. The kernel size corresponds to the 1D temporal convolutional layer used for processing input sequences. In each benchmark, the cross-modal attention and self-attention operations use an equal number of attention heads. Hyperparameter optimization is performed on the validation set.

**Table 1.**
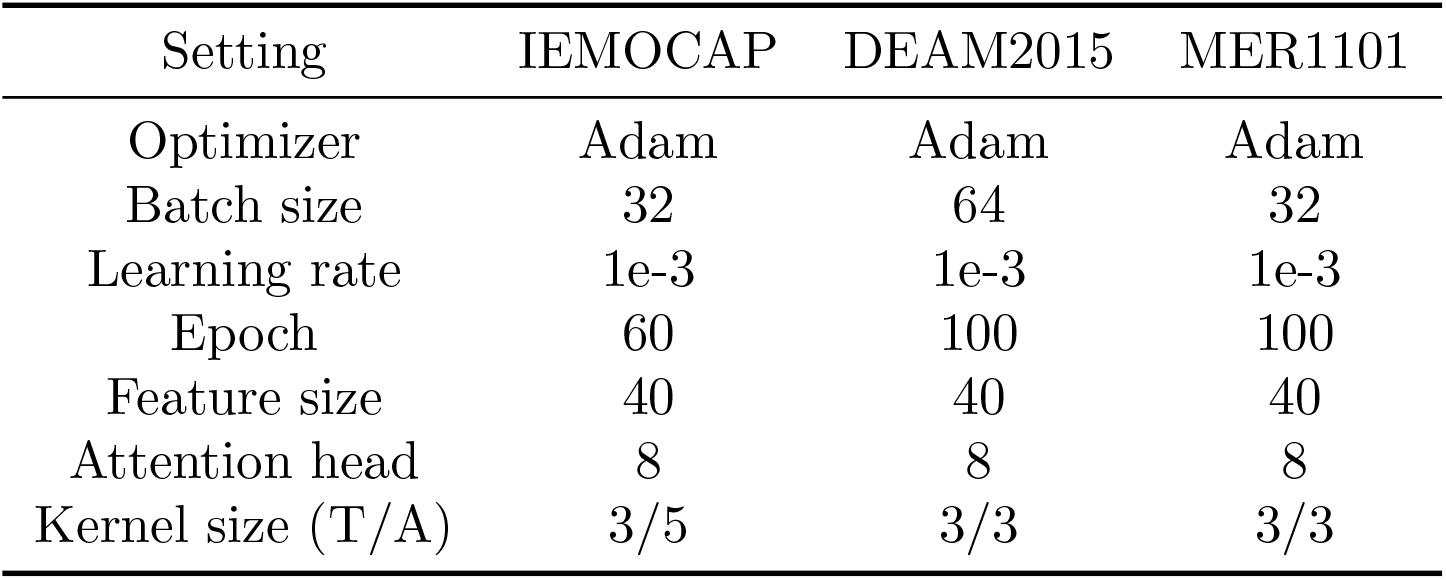
The hyperparameter settings adopted in each music emotion recognition benchmark.

| Setting | IEMOCAP | DEAM2015 | MER1101 |
| --- | --- | --- | --- |
| Optimizer | Adam | Adam | Adam |
| Batch size | 32 | 64 | 32 |
| Learning rate | 1e-3 | 1e-3 | 1e-3 |
| Epoch | 60 | 100 | 100 |
| Feature size | 40 | 40 | 40 |
| Attention head | 8 | 8 | 8 |
| Kernel size (T/A) | 3/5 | 3/3 | 3/3 |

### Performance Comparison

Our proposed approach is compared with several state-of-the-art baselines on IEMOCAP dataset with word-aligned setting, which visual and acoustic streams are manually aligned with textual words before fusion. As shown in Table 2, our approach demonstrates superior performance across different metrics compared to other baselines.

**Table 2.**
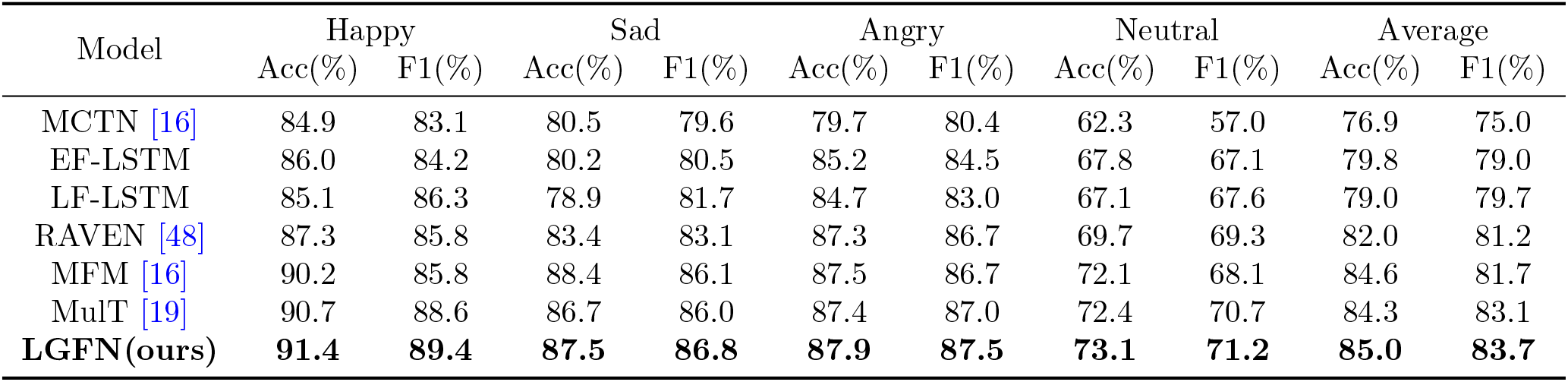
Comparison of other state-of-the-arts on the IEMOCAP benchmark. We report the the binary classification accuracy (Acc) and the F1-score on four emotion categories: happy, sad, angry and neutral.

We also compare our LGFN model with other MER methods from recent years on the DEAM2015 and MER1101 datasets. Notably, [34] uses the DEAM2014 dataset, which contains 744 songs, and [32–34] use RMSE for evaluation. Additionally, [9] converts numerical V-A values to binary representations for independent prediction every 0.5 seconds. All models are evaluated under identical experimental conditions, including the same datasets, evaluation metrics, and calculation methods. The experimental results are shown in Table 3. On the MER1101 dataset, our model outperforms others in all three metrics. On the DEAM2015 dataset, our model demonstrates strong recognition ability for arousal, though its performance in valence is slightly better than previous models. This might be due to less consistent valence annotations. We believe the predicted valence values on DEAM2015 are less reliable for DMER evaluation, as the C-CC values for valence from all models are near zero. The experiments show our model performs well on different datasets, especially in the arousal dimension. Overall, valence values in both datasets are more impoverished than arousal values, making valence prediction more challenging, a finding consistent with most previous studies.

**Table 3.** Comparison on the MER1101 and DEAM2015 dataset with other existing MER methods. We choose the CCC, PCC, and RMSE as evaluation metrics.

| Model | MER1101 dataset |  |  |  |  |  | DEAM2015 dataset |  |  |  |  |  |
| --- | --- | --- | --- | --- | --- | --- | --- | --- | --- | --- | --- | --- |
|  | Arousal |  |  | Valence |  |  | Arousal |  |  | Valence |  |  |
|  | CCC ↑ | PCC ↑ | RMSE ↓ | CCC ↑ | PCC ↑ | RMSE ↓ | CCC ↑ | PCC ↑ | RMSE ↓ | CCC ↑ | PCC ↑ | RMSE ↓ |
| CRNN [32] | 0.2798 | 0.5177 | 0.1625 | 0.0573 | 0.1033 | 0.2721 | 0.3488 | 0.5885 | **0.2197** | 0.0053 | -0.0292 | 0.3542 |
| BCRSN [9] | 0.1741 | 0.3770 | 0.3063 | 0.0660 | -0.0647 | 0.4143 | 0.3168 | 0.5148 | 0.2397 | 0.0125 | -0.0171 | 0.2914 |
| DNN [33] | 0.0529 | 0.0903 | 0.2372 | 0.0118 | 0.0017 | 0.2734 | 0.2757 | 0.4282 | 0.2483 | 0.0075 | 0.0031 | 0.3353 |
| MCRNN [34] | 0.0564 | 0.0918 | 0.2401 | 0.0155 | 0.0028 | 0.2752 | 0.2700 | 0.4396 | 0.2428 | 0.0137 | 0.0126 | 0.3135 |
| DAMFF [47] | 0.4223 | **0.6856** | 0.1494 | 0.1115 | 0.2004 | 0.2595 | 0.4203 | 0.6866 | 0.2401 | 0.0151 | 0.0366 | 0.3403 |
| **LGFN(ours)** | **0.4715** | 0.5923 | **0.1388** | **0.1348** | **0.3113** | **0.2309** | **0.4543** | **0.7022** | 0.2398 | **0.0160** | **0.0478** | **0.3310** |

### Ablation Studies

We conduct rigorous ablation studies across three benchmarks (MER1101, DEAM2015, IEMOCAP) to quantify the contribution of each core component. All experiments maintain identical configurations with the full LGFN model as the baseline. Performance is evaluated using CCC, PCC, and RMSE for regression tasks, while classification tasks employ single-trial accuracy (Acc_1_), five-trial average accuracy (Acc_5_), and F1-score.

### Necessity of Attentional Alignment Layer (ALL)

As shown in Table 4 and Table 5, the critical role of the Attentional Alignment Layer is demonstrated through systematic comparison with alternative fusion strategies. Replacing ALL with mean pooling, a method condensing the entire lyric sequence into a single static vector, reduces IEMOCAP Acc_1_ by 4.1%. Although this approach marginally outperforms complete ALL removal by preserving global semantic information, its inability to model temporal dynamics severely constrains fine-grained cross-modal interaction.

**Table 4.**
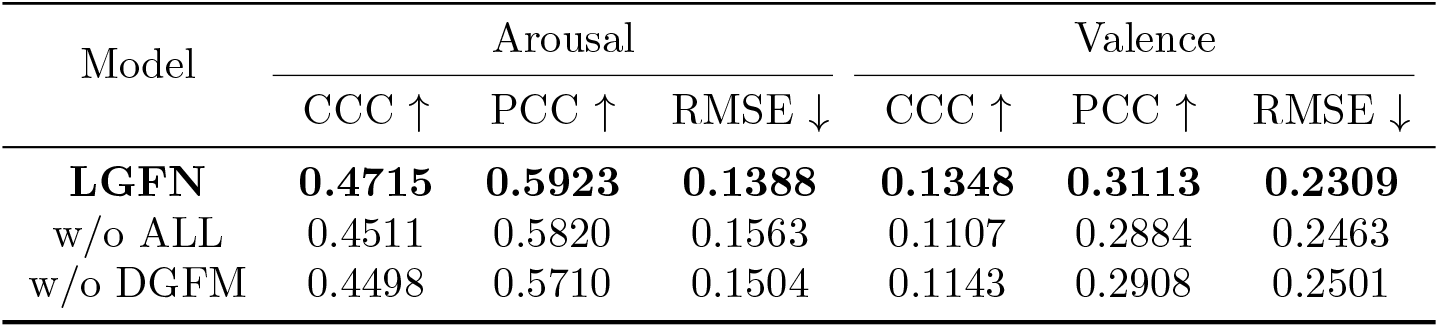
Ablation studies of arousal and valence on the MER1101 dataset.

**Table 5.** Ablation studies of arousal and valence on the DEAM2015 dataset.

| Model | Arousal |  |  | Valence |  |  |
| --- | --- | --- | --- | --- | --- | --- |
| | CCC ↑ | PCC ↑ | RMSE ↓ | CCC ↑ | PCC ↑ | RMSE ↓ |
| **LGFN** | **0.4543** | **0.7022** | **0.2398** | **0.0160** | **0.0478** | **0.3310** |
| w/o ALL | 0.4202 | 0.6866 | 0.2533 | 0.0124 | 0.0398 | 0.3366 |
| w/o DGFM | 0.4268 | 0.6720 | 0.2511 | 0.0136 | 0.0327 | 0.3421 |

Fixed-window alignment exhibits only limited improvement over mean pooling, yet still trails the full model by 3.7 accuracy points. This performance gap directly reflects its fundamental limitation: rigid segmentation mechanisms cannot adapt to natural lyric boundaries or musical phrase transitions.

Quantitative regression analyses reinforce these findings. ALL ablation degrades valence concordance correlation by 17.9% on MER1101 and 22.5% on DEAM2015. Such substantial declines underscore two inherent flaws in non-dynamic fusion: mean pooling discards essential sequential information crucial for frame-level emotion mapping, while fixed-window approaches force artificial alignments that misrepresent musical structure. The superiority of ALL stems from its content-aware dynamic alignment, which continuously adjusts lyric-audio correspondence through acoustic-linguistic feature interaction, enabling precise emotion recognition unattainable with coarse-grained fusion methods.

### Impact of Dynamic Gate Fusion Module (DGFM)

The adaptive nature of the Dynamic Gate Fusion Module is validated through systematic comparison with static fusion mechanisms. As demonstrated in Table 6, replacing DGFM with uniform static weighting, where audio and lyric features maintain fixed 0.5 contributions, reduces IEMOCAP Acc_1_ by 1.1% and Acc_5_ by 1.2%. This performance gap persists despite preserving the gate architecture, confirming that structural implementation alone cannot compensate for dynamic weighting capability. Regression analyses reveal parallel trends, with static fusion degrading arousal concordance correlation by 4.6% on MER1101 and 6.0% on DEAM2015. Crucially, the inferior performance of fixed-weight variants exceeds the margin between unimodal systems and full LGFN, demonstrating that context-aware reliability assessment surpasses manual parameter tuning. These findings establish that DGFM’s core innovation lies not in its gate structure, but in its real-time weight calibration based on feature credibility. In addition, as shown in Fig 3, our proposed methods can positively activate the music emotion recognition, which verify the effectiveness of the ALL and DGFM.

**Table 6.** Ablation study on the IEMOCAP benchmark. In each row, the corresponding component is progressively included into the model.

| Model design | Module |  | Acc<sub>1</sub>(%) | Acc<sub>5</sub>(%) | F1(%) |
| --- | --- | --- | --- | --- | --- |
|  | ALL | DGFM |  |  |  |
| LGFN (full model) | ✓ | ✓ | **84.8** | **85.0** | **83.7** |
| + Mean Pooling | ✗ | ✓ | 81.3 | 81.5 | 80.2 |
| + Fixed Window | ✗ | ✓ | 81.7 | 82.0 | 80.5 |
| + Audio-Only | ✓ | ✗ | 82.8 | 82.7 | 81.5 |
| + Lyrics-Only | ✓ | ✗ | 82.0 | 82.0 | 80.9 |
| + Fixed Gate (0.5) | ✓ | ✗ | 83.7 | 83.8 | 82.5 |
| + Fixed Gate (0.6) | ✓ | ✗ | 83.4 | 83.4 | 82.1 |

**Fig 3.**
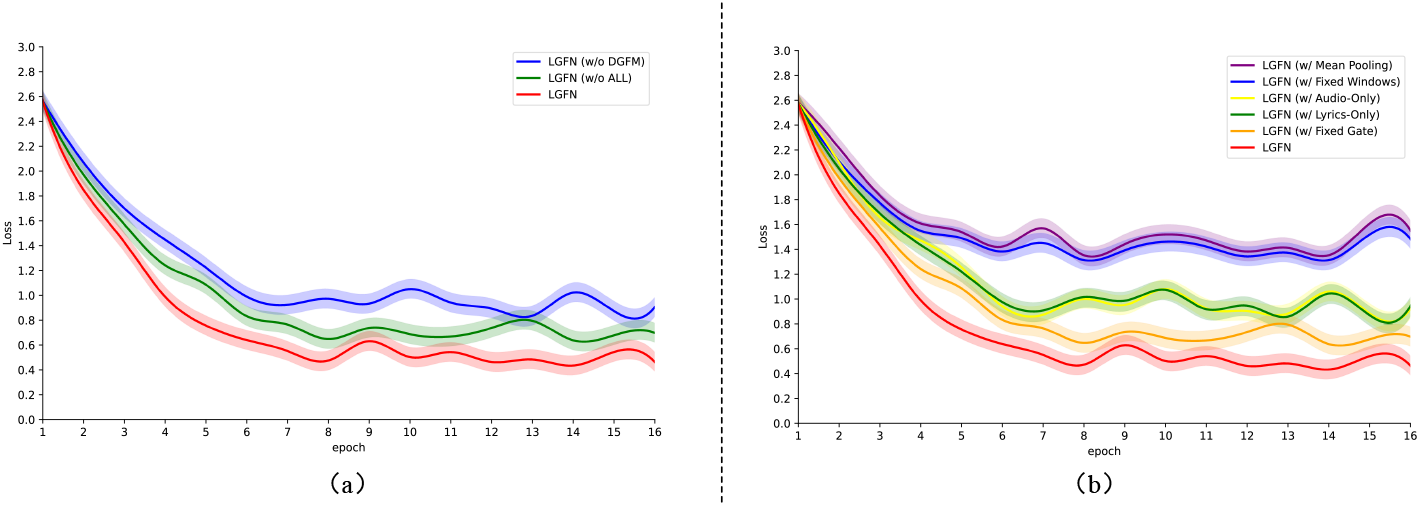
Comparison of emotion classification loss on music emotion benchmarks. (a) Comparison of emotion classification loss on the MER1101 dataset; (b) Comparison of emotion classification loss on the IEMOCAP dataset; w/o denotes without; w/denotes with.

### Multimodal Complementarity

Unimodal approaches exhibit inherent limitations that underscore cross-modal necessity. Refer to Table 6 and Fig 3, the Lyrics-Only configuration achieves 82.0% single-trial accuracy on IEMOCAP, while Audio-Only reaches 82.8%—both significantly trailing the full model’s 84.8%. This performance hierarchy persists across regression benchmarks, where valence correlation coefficients for unimodal systems consistently underperform multimodal integration by 12 ∼ 18%. Notably, simple feature concatenation without dynamic gating yields only marginal improvement over unimodal baselines, remaining 1.1 ∼ 1.4% below full LGFN. This critical gap confirms that mere modality combination is insufficient; effective emotion recognition requires DGFM’s reliability-aware fusion to resolve contradictory cues and amplify complementary signals during instrumental transitions or lyric-dense passages.

## Conclusion

This work proposes the Lyrics-Aware Gate Fusion Network LGFN, a novel framework addressing the dual challenges of audio feature ambiguity and lyrics-audio misalignment in music emotion recognition. To this end, we introduce a cross-modal attentional alignment layer that resolves temporal asynchrony through positional encoding, effectively synchronizing heterogeneous music streams. Moreover, a self-adaptive gating mechanism dynamically modulates audio-text feature contributions by suppressing semantically weak lyrics while amplifying emotionally salient words, enhancing discernment of nuanced emotions like excitement versus anger. The visualized gate weights further provide interpretable insights into modality reliability during fusion. Comprehensive evaluations confirm that our model establishes new state-of-the-art performance across diverse music emotion benchmarks.

